# Generation and Characterization of Autotetraploid Sweet Sorghum

**DOI:** 10.64898/2026.06.03.729885

**Authors:** Lizeth Dominguez Mendez, Brandon James, Wren Jenkins, Kankshita Swaminathan, Anthony J. Studer

**Affiliations:** Department of Crop Sciences, University of Illinois Urbana-Champaign, Urbana, IL 61801, USA; DOE Center for Advanced Bioenergy and Bioproducts Innovation, University of Illinois Urbana-Champaign; HudsonAlpha Institute for Biotechnology, 601 Genome Way, Huntsville, AL 35806, USA

**Keywords:** Autotetraploid, *Sorghum bicolor*, biofuels, polyploidy, sugar production

## Abstract

Increasing the diversity of biofuel crops can help meet energy demands while also stabilizing the domestic biofuel market. *Sorghum bicolor* is a promising feedstock for bioethanol production due to its sugar accumulation and storage in the stem in addition to its cellulosic biomass. Sorghum also exhibits high tolerance to abiotic stresses like extreme temperatures and drought. However, sorghum’s sugar production falls short when compared to current bioethanol feedstocks like maize and sugarcane. Therefore, to improve sorghum for the bioethanol market, an autotetraploid sorghum line was induced using colchicine treatments to increase cell size for greater sugar production and storage. Induced autotetraploid sorghum lines were validated with flow cytometry and screened using stomatal prints to detect larger stomatal cells. Two separate autotetraploid sorghum lines that were derived from the same M1 plant were characterized and evaluated for sugar production in a two-year field trial. The two autotetraploid lines displayed equal or improved performance when compared to their diploid equivalents for multiple juicing traits. Altogether, the data illustrate sorghum’s tolerance for autopolyploidy induction in an inbred background and suggest an opportunity for further improvements through progressive heterosis.

**SIGNIFICANCE STATEMENT:** Polyploidy has played a significant role in the improvement of some crop species. The characterization of a novel autotetraploid sweet sorghum line demonstrates the potential of increased sugar production in polyploids for biofuel applications.

## INTRODUCTION

It is predicted that by 2030 the demand for renewable energy in the transportation sector will increase by 50% (IEA, 2025). Thus, continued efforts to improve both current and future feedstocks are essential for meeting renewable energy demands. Sweet sorghum (*Sorghum bicolor* L. Moench) has been bred for its ability to produce sugar and store it in the stem (Regassa and Wortmann, 2014; Appiah-Nkansah *et al*., 2019). The sugars found in the stem of sweet sorghum can be fermented into ethanol for biofuels and bioproducts (Barcelos *et al*., 2016; Kasegn *et al*., 2024). Sorghum has potential utility as a biofuel feedstock because of its high tolerance to abiotic stressors such as elevated O_3_ concentrations, heat, drought, and salinity (Mathur *et al*., 2017; Prasad *et al*., 2021; Abuslima *et al*., 2022; Li *et al*., 2022; Frantová *et al*., 2024). When sorghum is compared to the leading bioenergy feedstocks, maize and sugarcane, sorghum requires less fertilizer input than maize and has shorter growing periods than sugarcane (Perrin *et al*., 2018; Eggleston *et al*., 2025). Despite these benefits, sorghum sugar yields are currently not competitive with those of either maize or sugarcane (Perrin *et al*., 2018; Eggleston *et al*., 2025). Thus, improving sugar yields in sorghum is necessary to increase its competitiveness within the bioenergy feedstock market.

Sorghum is a diploid species although it has close relatives that are polyploids (Price *et al*., 2005). Polyploidy is prevalent in crop species and could be implemented as a strategy to improve sugar yields of sorghum (Dominguez Mendez and Studer, 2025). Polyploid species are defined as containing two or more complete sets of homologous chromosomes and can generally be categorized into two broad categories, allopolyploids or autopolyploids (Tate *et al*., 2005). An allopolyploid arises through an outcross of two or more species in which the offspring contains full chromosome sets of each parent. Autopolyploids form when chromosomes from the same organism are doubled (Wendel and Doyle, 2005). Polyploids have been artificially induced to capitalize on increased cell size for greater yields (Comai, 2005).

The link between polyploidy and increased cell size has been documented for nearly a century and has been observed in several species (Müntzing, 1936; Dominguez Mendez and Studer, 2025). Crops, specifically alfalfa and potatoes, have long benefited from the increased yields associated with polyploidy and increased cell size (Cooper, 1939; Hancock Jr and Bringhurst, 1981). Additionally, leading cultivars of kiwifruit (*Actinidia chinensis* and *A. deliciosa*), strawberries (*Fragaria* x *ananassa*), and mint (*Mentha spicata*) are also polyploids (Wu *et al*., 2012; Bethke *et al*., 2022; Bharati *et al*., 2023). The model plant Arabidopsis has also been studied as both an auto- and allopolyploid where tetraploid varieties have increased flower and silique size along with a positive correlation between ploidy and guard cell size (Li *et al*., 2012; Corneillie *et al*., 2019). Similarly, inbred autotetraploid maize had larger stomata and pollen (Yao *et al*., 2011). Polyploid maize has been an invaluable resource furthering our understanding of gene dosage, heterosis, and inbreeding depression (Levings *et al*., 1967; Riddle and Birchler, 2008; Yao *et al*., 2011; Washburn *et al*., 2019; Batiru and Lübberstedt, 2024). The wide success of polyploidy species illuminate the potential application of an autopolyploid sorghum for increased sugar yields.

While larger cell size of polyploids has successfully increased yield in some species, higher ploidy is also associated with some drawbacks. For example, in species like Arabidopsis, maize, and alfalfa, polyploids had decreased seed set when compared to diploids (McCoy and Bingham, 1988; Li *et al*., 2012; Batiru and Lübberstedt, 2024). Other studies reported that tetraploids had a decreased growth rate, reduced fertility, and poor seed set (Kato and Birchler, 2006; Batiru and Lübberstedt, 2024). While reduced seed set can be limiting for production, past studies of newly synthesized autotetraploids have demonstrated that selection for increased seed set can overcome this drawback (Randolph, 1935; Luo *et al*., 1992). Additionally, polyploid bioenergy crops are not as impacted by reduced seed set because they are grown for their biomass or sugar accumulation. Thus, an autotetraploid sorghum has greater potential in the bioenergy sector where biomass and sugar are the harvestable products, not the grain.

Past literature suggests that increasing the ploidy level of sweet sorghum would increase the cell size and therefore sugar yield when compared to diploid sweet sorghum (Dominguez Mendez and Studer, 2025). This approach may be a strategy for making sorghum a competitive bioenergy crop. Various autotetraploid grain sorghums have previously been induced and described, but polyploidy has not been induced in a sweet sorghum background for sugar production (Doggett, 1964; Luo *et al*., 1992). If polyploidy is successful in increasing cell size in sorghum, we would expect an increase in sugar yield. To test this, an autotetraploid sweet sorghum (2n = 4x = 40) was induced through colchicine treatments and was characterized and evaluated for sugar production in a field setting.

## MATERIALS AND METHODS

### Plant Material

Sweet sorghum line ‘M 81E’ stock was used for tetraploid induction and untreated M 81E was used as the diploid control for all experiments. M 81E has been previously described in Broadhead *et al*., 1981 and is publicly available as accession PI 653411 through the Germplasm Resource Information Network (GRIN).

### Colchicine Treatments

Fifty seeds were sterilized with a 10% bleach solution and were placed in a 9cm petri dish with filter paper (Whatman Seed testing paper; 2181-090). Five seeds were placed on each plate, and water was added to wet the filter paper. The plates were sealed with parafilm and placed in an incubator set to 32°C to germinate in the dark for three days. When the coleoptiles were roughly 3cm in length, a 1mm section was removed from the tip using a single edge industrial razor blade (VWR: 55411-050). Each coleoptile was then inverted into a separate 1.5mL tube (Fisher Scientific 1.5ML MCT Graduated Natural: 05-408-129) containing a colchicine solution (Sigma-Aldrich:64-86-8) diluted to a 0.1% concentration with water. Special care was taken to ensure the meristem was fully submerged in the colchicine solution. Once all seedlings were inverted and submerged, a damp paper towel was gently placed over the roots to prevent drying. Seedlings were left in the 0.1% colchicine solution for 2.5 hours. After the treatment, seedlings were taken out and rinsed with water to remove the excess colchicine solution and immediately transplanted into a 50 well flat containing a soil mix previously described in (Twohey III *et al*., 2025). The flat was placed in a windowsill where the plants grew for 2 weeks under indirect sunlight. The two surviving plants were transplanted into 0.8L pots.

Then were again grown in the windowsill for an additional two weeks after transplanting. Flow cytometry was then used to analyze the DNA content of the two plants. Of the two colchicine treated plants, one plant was found to be a diploid and thus discarded, while the other colchicine treated plant was chimeric. 40 days after treatment, the chimeric M0 plant was then moved into a growth chamber, set to 12hr supplemental light, 32°C/24°C day/night temperatures until it was self-pollinated and harvested.

### Flow Cytometry

To screen potential autotetraploid plants, tissue was collected from both sides of the leaf midrib and placed on ice. Samples were chopped individually and also co-chopped with diploid control tissue. All samples were chopped using a single edge industrial razor blade (VWR: 55411-050) in 1mL of extraction buffer (425 mL water, 65 mL Hexylene glycol (2-methyl-2,4-pentanediol), 5mL 1M Tris (pH 8.0), 5mL 1M MgCl_2_). The chopped tissue solution was filtered using a 30µm cell strainer (Sysmex CellTrics; 04-004-2326) and placed on ice. Samples were then centrifuged at 500g to form a pellet and the supernatant was removed. Pellets were washed using 500µl of extraction buffer and were again centrifuged at 500g and the supernatant was removed. To stain nuclei, pellets were resuspended in 500µL of staining buffer (15mM HEPES, 1mM EDTA, 80mM KCl, 20mM NaCl, 300mM Sucrose, 1% Triton-X, 0.5mM Spermine, 0.5% PVP 40,000mw) containing a propidium iodide (PI; Sigma-Aldrich; 25535-16-4) solution. For every 1mL of staining buffer, the PI solution contained 100µL of PI and 10µg/mL of RNase, DNase-Free (Roche: 11119915001). Once resuspended, samples were incubated in the dark at room temperature for 1 hour before being placed back on ice. Flow cytometry was run using a BD LSR II Flow Cytometer (BD Bioscience) with voltages set to 555, 242, and 461 for forward scatter, side scatter, and PI respectively, and a BD FAC Symphony A1 cell analyzer (BD Bioscience) with voltages set to 319, 176, and 170 for forward scatter, side scatter, and PI respectively.

### Field Conditions

Field trials conducted in 2024 and 2025 were grown at the Crop Science Research and Education Center located in Urbana, Illinois, USA. 45 seeds were hand planted into 5m long rows with 1m alley and 0.8m spacing. In 2024, two rows of each M3 autotetraploid family were planted on June 13, 2024, and 4 rows of the diploid seed were planted 11 days later (June 24, 2024), to synchronize flowering times. In 2025, one row of each M3 family was planted on June 9, 2025, and again to synchronize flowering times, the diploid was planted 10 days later on June 20, 2025.

### Gas Exchange Measurements

Gas exchange measurements were taken in 2024 on the uppermost fully expanded leaf over 5 days starting at 71 days after planting (DAP) for the autotetraploids and 60 DAP for the diploid lines. Two portable photosynthesis systems (LI-COR 6800: Biosciences, Lincoln, NE, USA) were used for measurements and were alternated between sample groups. A total of 23 plants were measured, eight plants from M3 family B and diploid control plants and seven M3 family A plants. The Li-6800 chamber conditions were set to 28°C leaf temperature, pressure of 0.2 kPa, H_2_O concentration was controlled by relative humidity (RH) set to 60%, ambient, CO_2_ concentration (*C*_a_) controlled by the CO_2_ sample was set to 400µmol m^-1^, and light was set to 2000 µmol m^-2^ s^-1^ with 90% red and 10% blue light quality. Li-6800 chamber conditions were kept constant across all plants. Clamped leaves were given 30 minutes to acclimate, ensuring steady state was achieved before proceeding. Once at steady state, measurements were logged every 10 seconds for 5 minutes using an auto log program. At steady state, average CO_2_ assimilation (*A*_net_), stomatal conductance (*g*_s_), intracellular CO_2_ (*C*_i_), and intrinsic water use efficiency (i*WUE*) were calculated for each plant and treated as a single replicate and significance was determined using an ANOVA and Tukey’s post hoc test. i*WUE* was calculated with the equation (*A*_net_*/g*_s_).

CO_2_ response curves were measured immediately after steady state measurements were completed. The response curve was programmed to run through the following CO_2_ concentrations: 400, 300, 200, 100, 50, 75, 250, 400, 600, 800, 1000, 1200, 1500 µmol mol^-1^. Average *A*_net_, *g*_s_, *C*_i_, and i*WUE* for each sample group was calculated at each CO_2_ concentration point. Averages were used to plot an *A*/*C*_i_ curve with standard error bars shown. The maximum carboxylation capacity of phosphoenolpyruvate, *V*_pmax_, was calculated by extracting the initial slope of the *A*/*C*_i_ curve using the values of *C*_i_ < 50 µmol mol^-1^ (Von Caemmerer, 2000). The maximum CO_2_ saturated photosynthetic capacity, *V*_cmax_, was estimated using the horizontal asymptote of the *A*/*C*_i-_ curve by using a four-parameter non-rectangular hyperbolic function in SigmaPlot (Systat, San Jose, CA; Yendrek *et al*., 2017). Stomatal limitation was calculated using the equation described in (Long and Bernacchi, 2003).

### Stomatal Imaging

Autotetraploid plants grown in 2024 and 2025 were used to measure stomata size between ploidy levels. In 2024, stomatal prints were collected at 41 DAP for the autotetraploids and 30 DAP for the diploid and in 2025, prints were collected at 40 and 30 DAP respectively. A VistaVision Plain and Frosted Microscope Slide (VWR:16004-368) was prepped for each plant by labeling half the slide for the adaxial print and the other half for the abaxial print. During sample collection a marker was used to mark the adaxial side of the leaf, then a square was cut from the uppermost fully expanded leaf. To create stomatal prints, superglue was placed on one half of the slide to print the adaxial side then repeated for the abaxial side. Leaf samples were removed before the superglue fully dried, leaving stomatal prints behind. This was repeated on all autotetraploid M3 plants grown in the field. Slides were left overnight to completely dry. The following day, prints were imaged with an inverted OLYMPUS CK2 microscope using a 10x objective and a 3x MTV-3 microscope camera adapter. A total of six images were taken per sample, three of the adaxial side and three of the abaxial side. Images were then uploaded into BioDock AI (Biodock, AI Software Platform) where a pretrained AI model was used to count and measure images. Data was used to run an ANOVA and Tukey’s post hoc test to determine significant differences between diploid and autotetraploid sorghum.

### Stem Images and Diameter

In 2024, autotetraploid M3 stem samples were collected at anthesis, 110 DAP and 99 DAP for the autotetraploids and diploids, respectively. Plants were cut at the base, and the middle 7^th^ internode was removed, and stem diameter was recorded at the widest point. The 7^th^ internode was then placed on ice. Stem samples were vacuum infiltrated for 5-10 minutes in FAA solution (50mL EtOH, 5mL glacial acetic acid, 10mL formaldehyde, 35mL water) and incubated at room temperature for 16 hours. The FAA solution was then removed, and samples were vacuum infiltrated for 5-10 minutes with 50% ethanol then were left in the 50% ethanol for 1 hour. Wash steps were repeated with 60% and 70% ethanol. Samples were stored at 4°C in 70% ethanol until they were ready to cross section.

While holding the stems and observing the cuts with a SWIFT Stereo Microscope (S7-TP520-EA10-56), thin internode samples were hand sectioned into water using half of a wet Gillette Platinum razor blade. The internode sections were placed into a drop of water in the center of a Cellvis 35mm, #1.5 cover glass bottom dish (D35141.5N) and imaged using a Nikon AXR Confocal microscope with high-speed resonant scanner at 4× objective. Images were stitched and denoised using the Nikon Elements software.

Image analysis to count vascular bundles was performed using a UNET based convolutional neural network trained on a set of 160 images of Sorghum stem cross sections taken from the Sorghum Association Panel at Winfred Thomas Agricultural research station at Alabama A&M University (34.54’8”N 86.33’46” W, USDA hardiness zone 7B). This training was conducted by producing a set of 30 ground truth masks in which vascular bundles were marked with white dots over the entire area of the structure on a transparent image layer, which was then backed with a black layer to produce a binary mask. Images and masks were resized to 1792 by 1792 pixels, converted to 8 bit images, and thresholded on a scale from 0 to 255. These images were then separated into patches of 256 by 256 pixels, which were used to train the convolutional neural network.

Cross section segments of 2024 autotetraploid M3 sample stems were imaged using fluorescence microscopy, and these images were also put through the same modification steps - resizing, conversion to 8-bit, and thresholding. The model generated using the convolutional neural network was used to predict the presence of vascular bundles within these stems, and the number of predicted bundles within each stem was calculated.

### SLA and Isotope Leaf Collection

Leaf samples from the 2024 field trial were collected at 77 and 66 DAP for the autotetraploid and diploid respectively, to measure stable carbon isotope composition (δ^13^C, ‰). Leaf samples for δ^13^C were collected, weighed and folded as described in Twohey III et al., 2019. Samples were then sent to Washington State University Stable Isotope Core Laboratory for analysis using an elemental analyzer (ECS 4010, Costech Analytical, Valencia, CA) and a continuous flow isotope ratio mass spectrometer (Delta PlusXP, Thermofinnigan, Bremen; Brenna *et al*., 1997; Qi *et al*., 2003). Isotopic reference materials are interspersed with samples for calibration using two-point normalization to the Air-N_2_ and VPDB-LSVEC scales. Contribution of 17O is corrected by the IRMS software using the Santrock correction (Santrock *et al*., 1985).

Specific leaf area (SLA) samples were collected to determine ploidy effects to leaf thickness. To collect SLA samples, four 2.27cm^2^ leaf punches were collected from 10 individual plants per group. After collection, samples were placed in an incubator set to 65°C to dry. Once fully dried, each leaf punch was weighed individually then averaged per plant to calculate SLA values for each plant. Averages were used to run ANOVA and Tukey’s post hoc test.

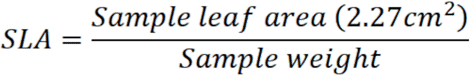

### Field Phenotyping

In 2024, plant height and total leaf number were measured on 10 plants from each autotetraploid family and a diploid control at anthesis. Plant height was measured as the distance between the base of the main stalk and the node of the flag leaf. Total leaf number was measured by counting the leaves of the main stalk, including the flag leaf.

Sorghum juicing was conducted 30 days after anthesis. In 2024, autotetraploids reached anthesis at 110 DAP and the diploids reached anthesis 99 DAP, and in 2025 autotetraploids and diploids reached anthesis at 111 and 101 DAP. Therefore, they were juiced at 140 and 129 DAP in 2024, and at 141 and 131 DAP in 2025. On the day of juicing, sorghum plants were cut down at the base of the stalk including all tillers. Plants were grouped into 5 plant bundles, and each bundle was treated as one replicate. A total of eight bundles were collected for each ploidy group in 2024, and four bundles were collected for each ploidy group in 2025. After all bundles were collected, the total fresh weight of the stalks and panicles were recorded for each bundle.

Panicles were removed before juicing. Bundles were juiced using a 2013 Edwards Engineering Corp Stationary Sugarcane Juice Extraction Unit (SCM-APL-3C, Edwards Engineering). Total juice volume, bagasse weight, and °brix were measured. A Palm Abbe digital refractometer (MISCO: PA-202) with a 200mL of juice sample was used to measure °brix. Fresh bagasse and panicles were placed in 65°C dryers to dry. After one-week, dry weights were collected on both the bagasse and panicles. Total dry weight does not include seed weight. The following equations were used to determine juicing ability.

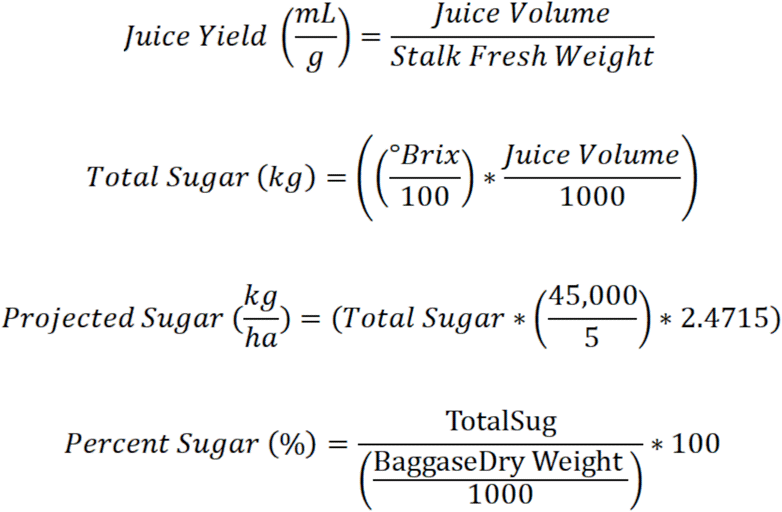

### Statistics

Using RStudio version 2025.5.1.513, the best fit model was found for stomatal prints and juicing traits (Posit Team, 2025). For stomatal traits, year was found to have no significant effect, and the following linear model was used for statistical analysis.

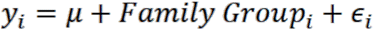

Where *y_i_* is the stomatal size response variable for the *i^th^* family group (Family A, Family B, or diploid), and ∈_i_ is the residual error. For juicing traits, a mixed model was used using the lme4 package (Bates et al., 2015). For juicing traits, year was found to be significant and was included in the following model:

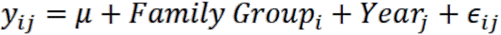

Where *y_ij_* is the juicing response variable for the *i^th^* family group in the *j^th^* year, where family group is a fixed variable and year is a random variable.

## RESULTS

### Generating Autotetraploid Sorghum

Flow cytometry was used to quantify DNA content to distinguish diploid individuals from autotetraploids. Of the two M0 plants that were analyzed with flow cytometry, one showed 2C and 4C peaks comparable to the diploid control (Figure S1A-B). The 2C peak represents the G1 phase of the diploid cells and the 4C peak represents the G2 phase of the diploid cells. The other M0 plant was chimeric, with 2C and 4C peaks comparable to the diploid control, but also a 8C peak indicating the plant possessed cells that were 4x (Figure S1C). In this case the 4C peak represented both diploid cells in G2 and autotetraploid cells in G1 phase of the cell cycle, while the 8C peak represented the G2 phase of the autotetraploid cells. The chimeric M0 plant was self-pollinated, resulting in M1 seed. M1 plants were grown and tested using flow cytometry. The M1 samples were analyzed with flow cytometry individually, and as co-chopped samples with equal quantities of diploid tissue to identify true autotetraploids with three distinct peaks. M1 flow cytometry results confirmed an autotetraploid sorghum plant (Figure 1). In the known diploid control of the same genotype, a 2C peak was shown at 48,000 PI-A, and a 4C peak at 96,000 PI-A (Figure 1C). Based off the diploid, expected peak values for an autotetraploid individual would be 96,000 PI-A for the 2C peak and 192,000 PI-A for the 4C peak, which was similar to the observed values of the M1 autotetraploid, 91,000 and 180,000 PI-A for the 2C and 4C peak respectively (Figure 1A). A co-chopped sample was also run to further confirm the ploidy of our autotetraploid where three C peaks are observed at 39,000, 79,000, and 159,000 PI-A respectively (Figure 1B). The M2 and M3 generations of autotetraploids were also validated using flow cytometry demonstrating the stable inheritance of the autotetraploid (Figure S2 and S3). Family A and B were derived from selfing individual M2 plants. Plants were selected based on vigor and seed set.

**Figure 1.**
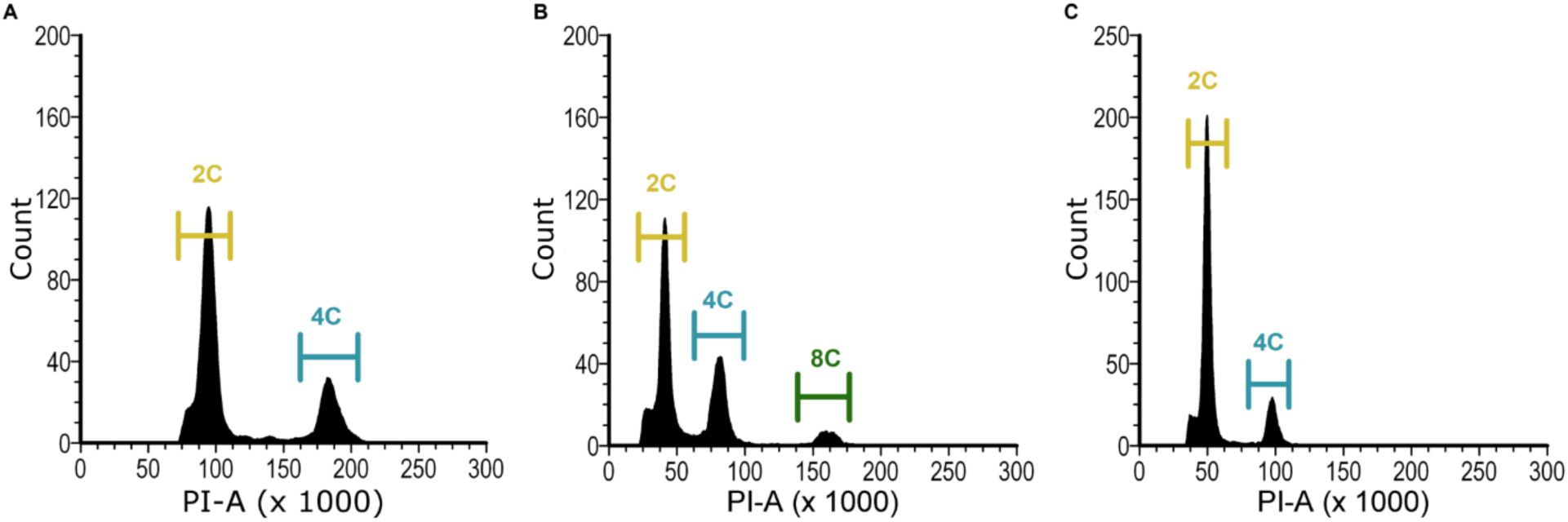
Flow cytometry results of (A) the 4x M1 sample alone, (B) co-chopped sample of the diploid and 4x M1 plant, and (C) the diploid sample alone.

### Cell Size is Positively Correlated With Ploidy

To determine if an increase in ploidy resulted in larger cells in autotetraploid sorghum, stomata and parenchyma cells were imaged and compared to diploid controls of the same genetic background. Stomata and parenchyma cells were measured in M3 plants from Family A and B. Stomata length, width, and area were significantly larger in both autotetraploid families on both the abaxial and adaxial leaf surfaces (Figure 2A, Figure S4 A-B), while stomata density significantly decreased (Figure 2B). Autotetraploid sorghum had larger, but fewer stomata compared to diploid sorghum. Similarly, when observing parenchyma cells in sorghum stems, autotetraploid plants also had larger but fewer parenchyma cells compared to diploid sorghum (Figure 2C-D). Thus, larger cells were observed as a result of increased ploidy level across multiple tissues. However, when comparing sorghum stem diameter, there were no differences across groups (Table 1). When counting the number of vascular bundles (VB) in the stem, with diploid sorghum having more VB than M3 family B, but not more than M3 family A (Table 2).

**Figure 2:**
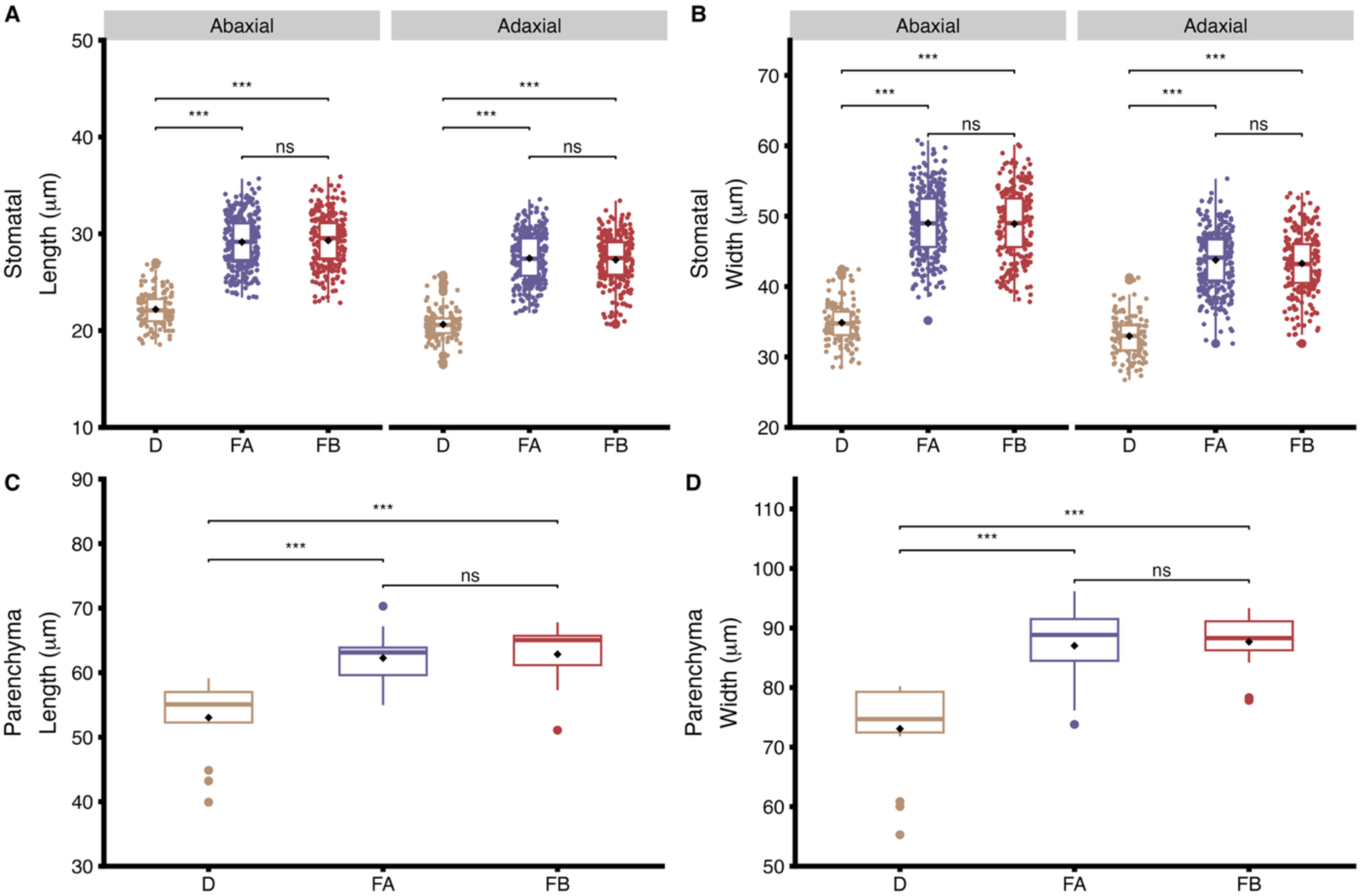
Box plots showing (A) Stomatal area of individual stomatal averaged per plant, (B) Stomatal density (0.66mm^2^) on both the abaxial and adaxial side of the leaves, (C) Parenchyma area of individual cells averaged per plant shown, and (D) Parenchyma density from whole stem cross sections. *p*-value < .001 denoted with ***. Significance was determined using ANOVA and Tukey’s post hoc test.

**Table 1:**
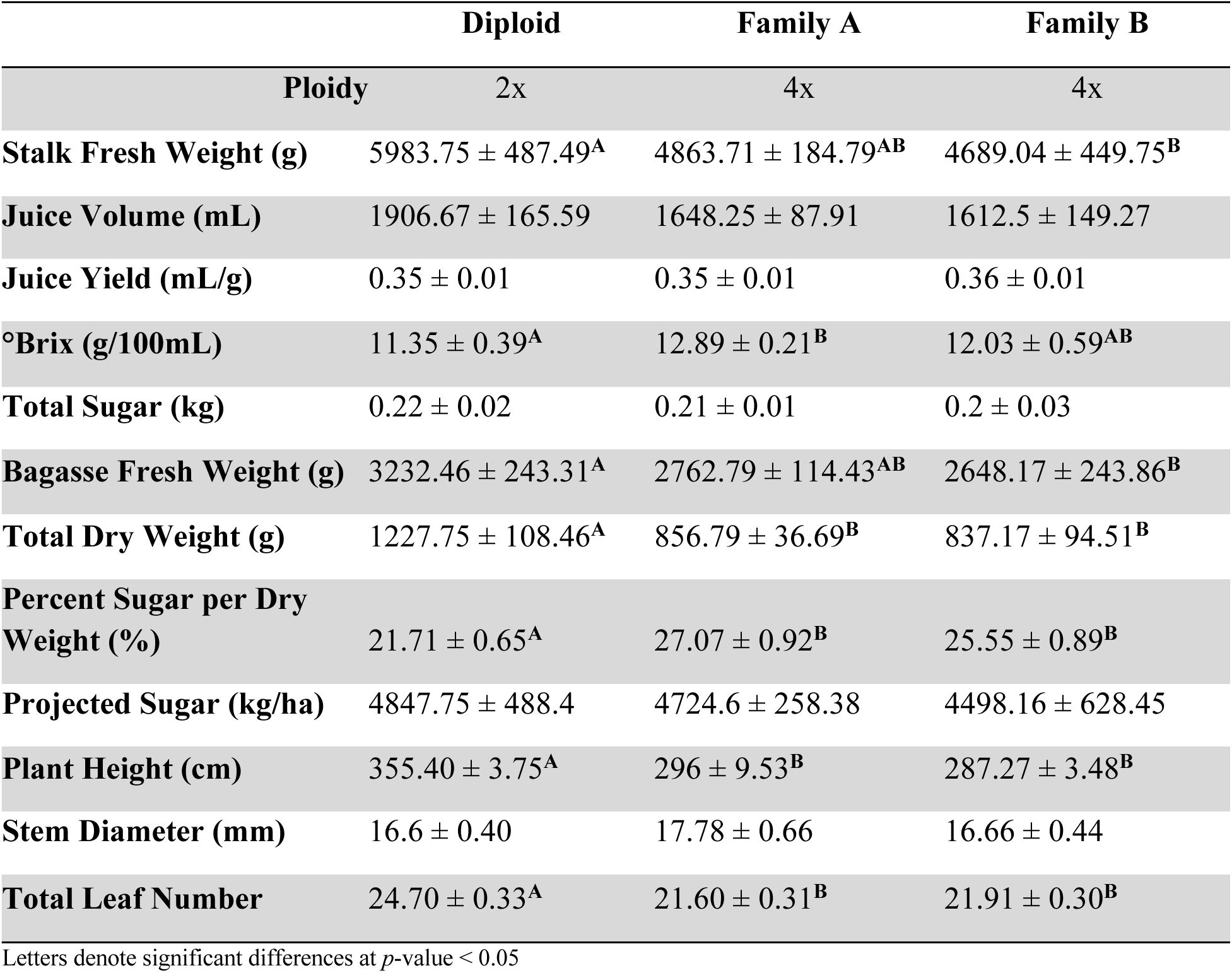
Juicing and growth traits for autotetraploid families and the diploid control. Significant differences (ANOVA & Tukey’s post hoc) denoted with letters to show grouping. No letters indicate no differences across any grouping. *n* = 8 for juicing measurements, and *n* =10 for growth traits.

**Table 2:** Steady-state measurements and leaf traits for autotetraploid families and the diploid control. Significant differences denoted with letters to show grouping. No letters indicate no statistical differences across any grouping. For steady state measurements *n* = 8 for the diploid and autotetraploid family B group and *n* = 7 for the autotetraploid family A group. For all leaf traits *n* = 10 for all ploidy groups.

|  | Diploid | Family A | Family B |
| --- | --- | --- | --- |
| Ploidy | 2x | 4x | 4x |
| **$A_{\text{net}}$ ($\mu\text{mol m}^{-2} \text{s}^{-1}$)** | $45.73 \pm 1.33^{\text{A}}$ | $39.07 \pm 1.57^{\text{B}}$ | $37 \pm 1.1^{\text{B}}$ |
| **$C_i$ ($\mu\text{mol m}^{-1}$)** | $110.03 \pm 6.04$ | $128.12 \pm 5.6$ | $130.57 \pm 8.01$ |
| **$g_s$ ($\text{mol m}^{-2} \text{s}^{-1}$)** | $0.27 \pm 0.013$ | $0.24 \pm 0.01$ | $0.23 \pm 0.013$ |
| **$iWUE$ ($\mu\text{mol mol}^{-1}$)** | $171.6 \pm 3.92$ | $160.97 \pm 3.51$ | $159.66 \pm 5.1$ |
| **Percent Nitrogen** | $3.12 \pm 0.14$ | $3.11 \pm 0.12$ | $3.14 \pm 0.11$ |
| **leaf $\delta^{13}\text{C}$ ‰** | $-12.57 \pm 0.04$ | $-12.6 \pm 0.05$ | $-12.56 \pm 0.08$ |
| **Percent Carbon** | $43.1 \pm 1.46$ | $43.51 \pm 1.37$ | $43.93 \pm 0.7$ |
| **SLA** | $234.71 \pm 6.95^{\text{A}}$ | $210.69 \pm 4.99^{\text{B}}$ | $224.77 \pm 4.38^{\text{AB}}$ |
| **Number of Vascular Bundles** | $338.0 \pm 8.17^{\text{A}}$ | $321.2 \pm 4.50^{\text{AB}}$ | $296.4 \pm 8.38^{\text{B}}$ |

Letters denote significant differences at $p$ -value $< 0.05$

### Assimilation Rates Do Not Increase With Ploidy

Gas exchange measurements were conducted to determine if the larger stomatal cells influenced photosynthetic properties in autotetraploid sorghum. Steady-state gas exchange measurements were used to compare *A*_net_, *g*_s_, *C*_i_, and i*WUE* between ploidy groups. No significant differences were observed for *C*_i_, *g*_s_, and i*WUE* when comparing diploid and autotetraploid plants (Table 2). However, both steady-state and *A*/*C*_i_ curve measurements showed diploid plants had significantly higher *A*_net_ when compared to either autotetraploid family (Table 2). The CO_2_ response curves revealed that both autotetraploid sorghum families had significantly lower *V*_cmax_ than the diploid control, but the two autotetraploid families were not significantly different from each other (Figure 3). Similar observations were seen for *V*_pmax_ (Figure 3).

**Figure 3:**
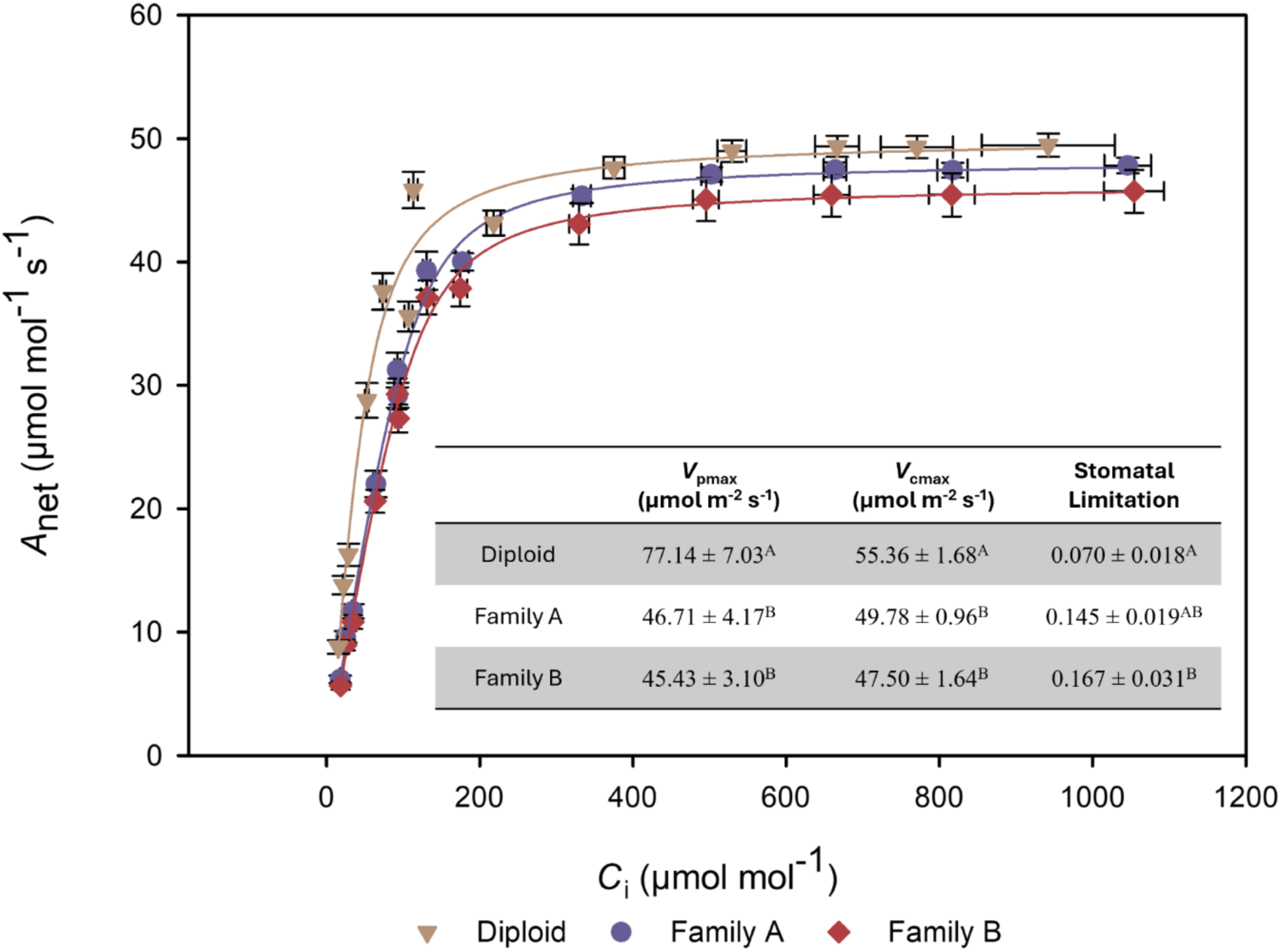
*A*/*C*_i_ curve with standard error bars for both autotetraploid families and diploid control. Embedded table shows respective values for both *V*_pmax_ and *V*_cmax_. n = 8 for Family B and the diploid and n=7 for Family A. Significant differences (ANOVA & Tukey’s post hoc) denoted with letters.

Autotetraploid plants were also found to have increased stomatal limitation to *A*_net_ (Figure 3). Autotetraploid family B was found to have significantly increased stomatal limitation when compared to the diploid control. Meanwhile, autotetraploid family A was not significantly different than either autotetraploid family B or the diploid, however it did appear to trend higher than the diploid.

### Growth Observation in Autotetraploid Sorghum

Higher ploidy levels have been shown to decrease growth rates in plant species (Riddle and Birchler, 2006; Li *et al*., 2012). Autotetraploid families A and B, along with diploid sorghum, were phenotyped for plant height, flowering time, and total leaf number to determine if increased ploidy altered sorghum growth. Both autotetraploid M3 families flowered later than the diploid. Plant height and leaf number were reduced in both autotetraploid families when compared to diploid sorghum but no differences between autotetraploid families were observed (Table 1). Leaf tissue was collected to measure carbon isotope composition (δ^13^C), carbon and nitrogen content, and SLA. There were no significant differences across ploidy groups when comparing δ^13^C, percent carbon, or percent nitrogen (Table 2). However, when comparing SLA values, M3 family B displayed an intermediate phenotype, as it was not significantly different from either M3 family A or the diploid, but the diploid did have significantly higher SLA values than M3 family A (Table 2).

### Autotetraploid Sorghum Produced More or Equal Amounts of Sugar

Juicing traits were collected to determine juicing abilities of autotetraploid sorghum and to determine if the reduction in *A*_net_ altered sugar production. A total of 12 bundles were juiced from each autotetraploid family and the diploid control across two years. When comparing juice volume, juice yield, total sugar, projected sugar, and stem diameter there are no significant differences across all groups (Table 1). However, when comparing stalk fresh weight and bagasse fresh weight of the diploid and autotetraploid sorghum families, the diploid was significantly higher than M3 family B, but not significantly different than M3 family A, nor were there significant differences between the two autotetraploid families (Table 1). Additionally, when comparing total dry weight, the diploid had significantly more biomass than both autotetraploid families (Table 1). Thus, to account for differences in biomass, samples were normalized to dry weight and percent sugar per dry weight was calculated. Interestingly, both autotetraploid families had significantly higher percent sugar per dry weight than the diploid control when normalizing for growth differences (Table 1). Additionally, M3 family A outperformed the diploid in °brix, or sugar concentration, but there were no significant differences between the diploid and M3 family B, nor were there significant differences between the two autotetraploids (Table 1). Despite both autotetraploid families having slower growth rates and reduced *A*_net_, autotetraploid family B was able to maintain, while family A was able to increase °brix, which resulted in both families having an increased percent sugar per dry weight (Table 1). Together, these data suggests that sorghum can tolerate the polyploid event as an inbred.

## DISCUSSION

Polyploid sorghum holds great promise for reaping the potential benefits associated with higher ploidy (Dominguez Mendez and Studer, 2025). Consistent with previous reports, colchicine treatments were successful in inducing an autotetraploid sorghum (Doggett, 1964; Luo *et al*., 1992). While flow cytometry can be used to confirm ploidy level, we have also shown that stomatal prints were an effective method of rapidly screening for autotetraploid sorghum plants (Figure 2). Advancing autotetraploid sorghum lines to the M3 generation demonstrated the stable inheritance of the polyploidy.

While increasing the ploidy level of sorghum led to larger stomata and parenchyma cells, there was a corresponding reduction in cell density along with no change in stem diameter (Figure 2; Table 1). Similar findings have been reported in induced polyploid species such as, maize, *M. x giganteus*, Arabidopsis, and mint (Riddle *et al*., 2006; Yu *et al*., 2009; Yao *et al*., 2011; Corneillie *et al*., 2019; Bomblies, 2020; Bharati *et al*., 2023). The reduction in cell density can be attributed to the reduced number of cell divisions observed in polyploid species (Tsukaya, 2008; Corneillie *et al*., 2019). Higher ploidy cells require more energy to complete a round of cell division, which reduces the total number of cell divisions compared to diploid species (Tsukaya, 2008). Past studies have also observed the trade-off of larger but fewer cells to maintain organ size; however, the underlying cause of this trade-off remains unknown (Robinson *et al*., 2018).

Larger cells have the potential to alter numerous cellular processes including those related to photosynthesis and gas exchange physiology. For example, the observed decrease in *A*_net_ in autotetraploids could be due to larger cells having decreased rates of diffusion of photosynthates from mesophyll cells into bundle sheath cells. A previous study showed that increased ploidy was strongly correlated with bundle sheath cell size but not mesophyll cell size, which would rule out a diffusion limitation (Warner and Edwards, 1989). Although the exact size of mesophyll and bundle sheath cells were not measured here, the presence of the carbon concentrating mechanism in C_4_ species would likely result in diffusion rate differences being negligible. Larger stomatal cells also result in larger pore apertures which can decrease i*WUE* (Lawson and Blatt, 2014; Lawson and Vialet-Chabrand, 2019). Thus, because the autotetraploids had larger but fewer stomata, this led to questions about the effects of autotetraploid stomata on gas exchange. The larger but fewer stomata offset each other leading to no significant differences to overall *g*_s_ or i*WUE*. Although, both *g*_s_ and i*WUE* trended lower in both autotetraploid M3 families (Table 2). A previous attempt to increase i*WUE* in diploid sorghum through reduced stomatal density was offset by an increase in stomatal size (Lunn *et al*., 2024). Furthermore, studies suggest that severely decreasing stomatal density can limit CO_2_ diffusion into the leaf, which reduces *A*_net_ (Ferguson *et al*., 2024). In this study, although autotetraploids at steady-state conditions had significantly lower *A*_net_, there were no significant differences in *C*_i_ across all groups, indicating that the reduction in *A*_net_ was not due to a CO_2_ limitation (Table 2 & Figure 3).

Stomatal limitation was also calculated from *A*/*C*_i_ curves to determine if the reduced *A*_net_ was due to stomatal response to dynamic conditions. Interestingly, autotetraploid family B had a significantly greater stomatal limitation, while family A had an intermediate phenotype, and was not significantly different than either family B or the diploid (Figure 3). Regardless, both autotetraploids trended, or were significantly limited by stomata under dynamic conditions when compared to the diploid. Thus, the reduced *A*_net_ observed here may be due to the larger stomata limiting photosynthesis under changing environmental conditions. Also supporting the increased stomatal limitation in autotetraploid sorghum is that during *A*/*C*_i_ curves, autotetraploid M3 families maintained relatively low *g*_s_ values throughout the entire curve, even at low *C*_i_ concentrations (Figure S5). In contrast, diploid sorghum responded to low *C*_i_ by increasing *g*_s_ for greater CO_2_ uptake (Figure S5). The phenomenon of larger but slower stomata has been previously shown and attributed to increased amounts of solutes needed to trigger a stomatal response (Drake *et al*., 2013; Lawson and Blatt, 2014). In addition, past studies on diploid sorghum have shown a correlation between stomatal size and delayed stomatal closure (Al-Salman *et al*., 2023). Thus, the decreased stomatal response rate in autotetraploid sorghum may be driving the stomatal limitation and therefore, also reduced *A*_net_.

Both autotetraploid M3 families had significantly lower *A*_net,_ *V*_cmax_, and *V*_pmax_ (Figure 3). *V*_cmax_ represents the maximum carboxylation rate and is used as a proxy for rubisco activity; while *V*_pmax_ represents the carboxylation capacity of phosphoenolpyruvate (PEPc) and can serve as a proxy for PEPc activity (Long and Bernacchi, 2003). Thus, because both *V*_cmax_ and *V*_pmax_ of the autotetraploids were significantly lower than the diploid, perhaps enzyme activity does not scale proportionally with cell size in polyploids. In this study we did not conduct enzyme activity assays, which could be used in the future to determine the cause of decreased *V*_pmax_ and *V*_cmax._ Interestingly, although autotetraploid sorghum displayed decreased *A*_net_, equal, if not greater sugar concentration in the stem was observed (Table 1).

The aim of this study was to determine if increasing the ploidy level of sorghum was a viable strategy for increasing sugar production through increased cell size. Autotetraploid sorghum was able to maintain sugar production and storage abilities. Both autotetraploid M3 families were stable and demonstrated that sorghum could tolerate autopolyploidy in an inbred state. M3 family A was equal to, if not better than, the diploid control for °brix, stalk fresh weight, and percent sugar per dry weight (Table 1). In contrast, family B showed variability when compared to the diploid for the same traits (Table 1). These results suggest that genome stabilization after a whole genome duplication event may be variable in early generations of offspring given that the autotetraploid families have identical genetic backgrounds. Future studies may investigate polyploidization effects between the two lines to elucidate the phenotypic differences observed here.

In this study, autotetraploid sorghum as an inbred did not demonstrate negative impacts on sugar production in response to a higher ploidy level. The same cannot be said for inbred autopolyploids in other species. For example, inbred autotetraploid maize had decreased ear length, reduced fertility, and reduced seed set (Riddle *et al*., 2006; Yao *et al*., 2011; Batiru and Lübberstedt, 2024). Despite the autotetraploid inbreds having reduced fitness, hybrid autotetraploid maize had increased heterosis compared to its diploid counterpart across multiple traits (Levings *et al*., 1967; Riddle and Birchler, 2008; Washburn *et al*., 2019). Notably, double cross autotetraploid maize hybrids exhibited progressive heterosis which further increased the amount of heterosis observed (Levings *et al*., 1967; Washburn *et al*., 2019). Progressive heterosis is the increase in hybrid vigor observed in a double cross hybrid when compared to parental inbreds and single cross hybrids (Groose *et al*., 1989). Given that hybrid autotetraploid maize displayed increased vigor through progressive heterosis regardless of its poor tolerance to polyploidy as an inbred, it is likely that progressive heterosis can be used to further increase autotetraploid sorghum sugar yields.

## CONCLUSION

In this study, an autotetraploid sweet sorghum line was developed in an effort to increase sugar production. Increasing the ploidy level of sorghum resulted in increased cell size, specifically larger stomata and parenchyma cells. However, cell density of stomata and parenchyma cells decreased in the autotetraploids. Despite no observable differences in cell size between autotetraploids, sugar yield varied between the two autotetraploid sorghum lines.

Additionally, gas exchange measurements indicated that the autotetraploids had decreased photosynthetic parameters but maintained sugar yields compared to the diploid control.

## ACKNOWLEDGMENTS

Thank you to my labmates who helped me conduct the juicing trials for this study, Bryan Warsaw for his help managing field trials at Illinois, James Schnable for the sorghum association panel (SAP) seeds, and Xianyan Kuang for managing the SAP field at Alabama A&M University. We’d like to thank Daija Stepney for developing the first version of the machine learning pipeline to count vascular bundles, Steve Moose for thoughtful discussions about sorghum, and George Hodnett for early conversation and guidance on flow cytometry.

## SUPPLEMENTAL FIGURES AND GRAPHS

**Figure S1:**
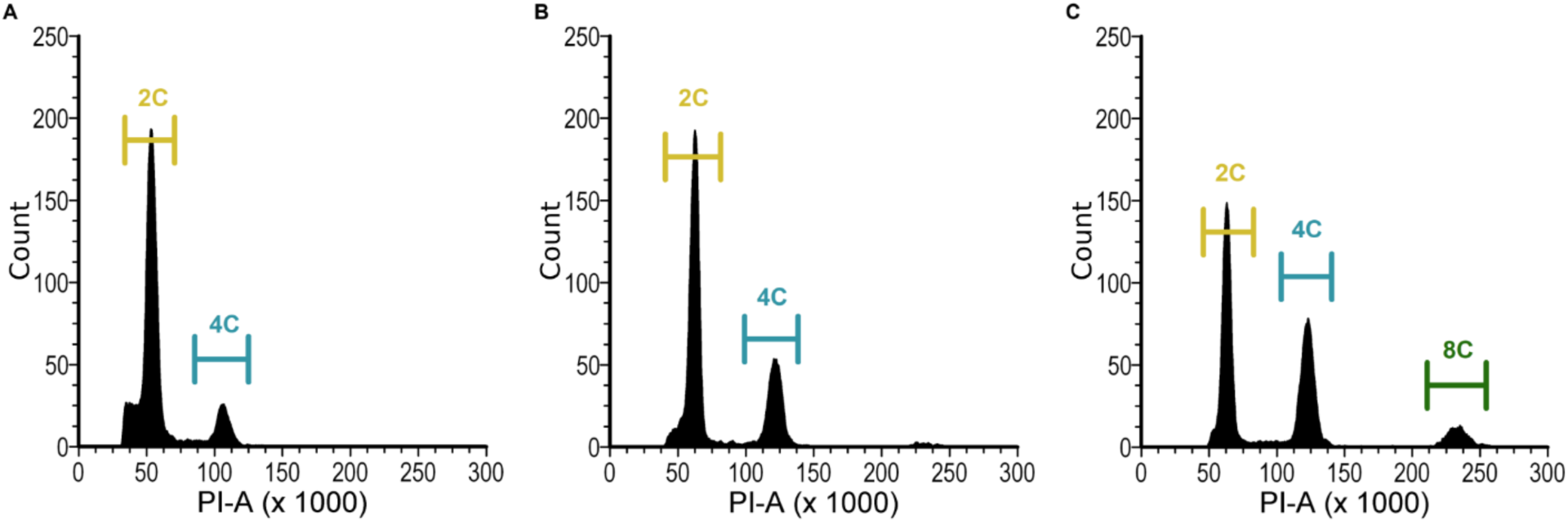
Flow cytometry results of the (A), diploid control, (B) M0 plant that was not affected by colchicine treatments and the (C) M0 plant that was affected by colchicine treatments and was chimeric.

**Figure S2:**
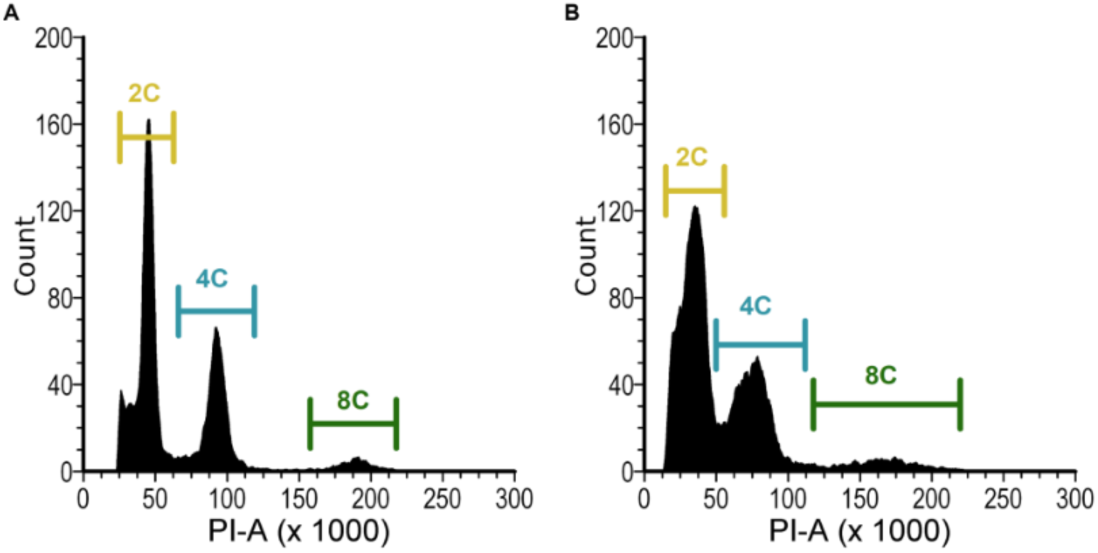
Flow cytometry results from the M2 generation for “A” and “B” families. Panel (A) are results for the family A M2, and panel (B) shows the results for the family B M2.

**Figure S3:**
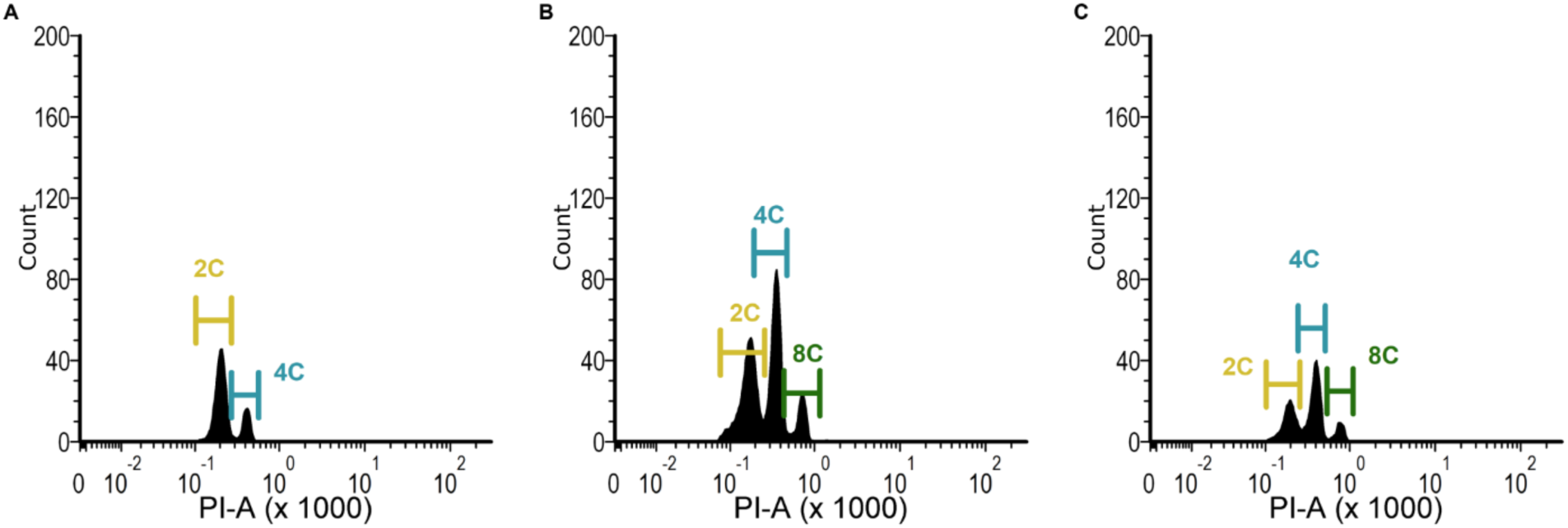
Flow cytometry results using A1 Symphony cytometer. (A) Diploid control, (B) M3 Family A representative, and (C) M3 Family B representative.

**Figure S4:**
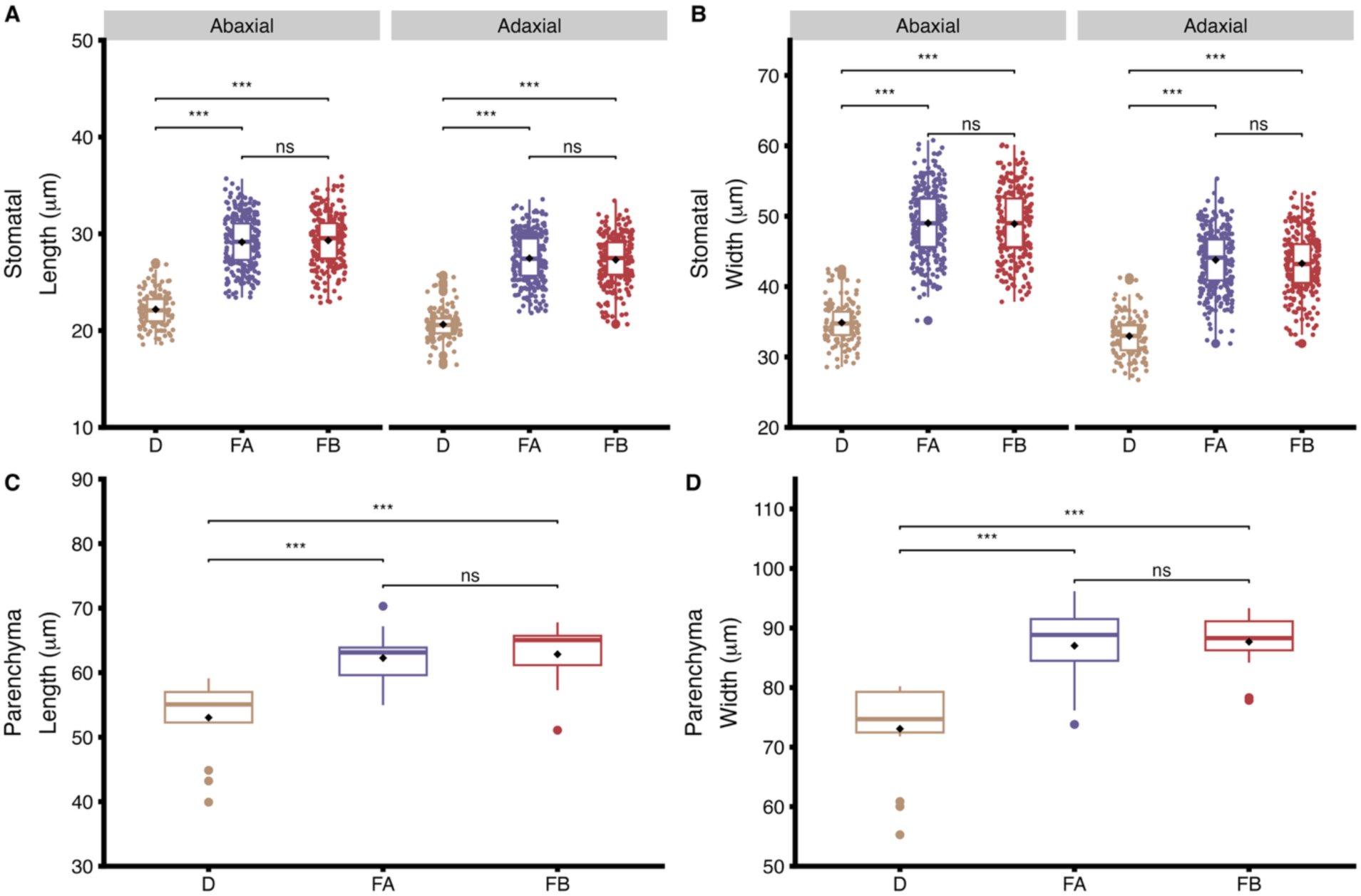
Boxplots showing Stomatal (A) length, (B) width, and parenchyma (C) Length and (D) width in number of pixels. *p*-value < .001 denoted with ***. Significance was determined using ANOVA and Tukey’s post hoc test.

**Figure S5:**
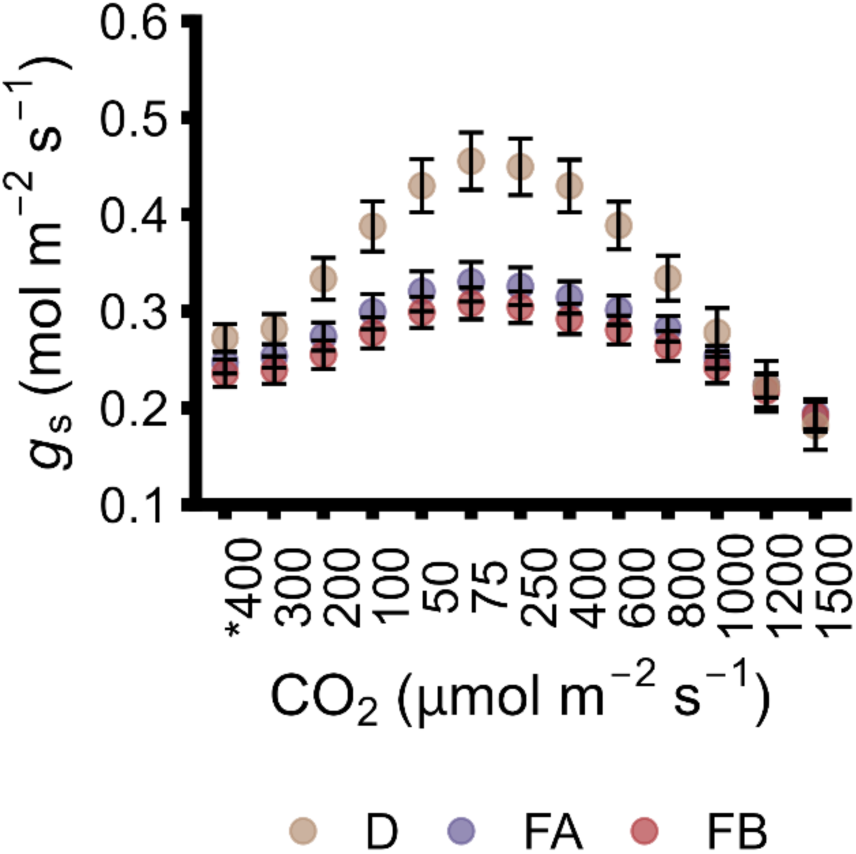
*g*_s_ measurements throughout the CO2 response curve shown for both autotetraploid families and diploid control. *400 is the first CO_2_ set point of the response curve.

## Tables

**Table S1:**
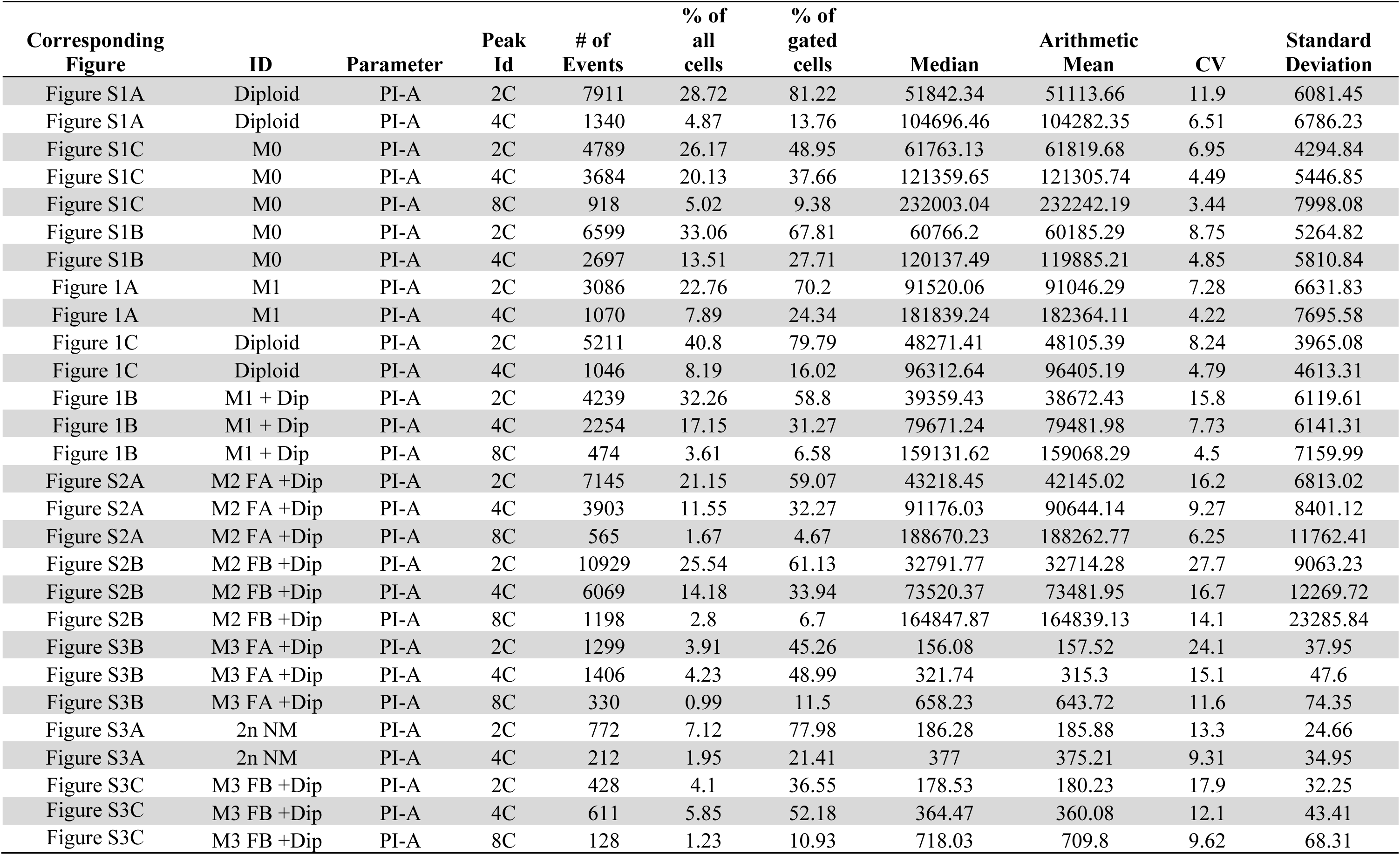
Statistics from the flow cytometer histograms shown.

